# Intergenerational transfer and sex differences of DNA methylation patterns in the Pacific oyster (*Crassostrea gigas*)

**DOI:** 10.1101/2022.02.22.481396

**Authors:** Yongguo Li, Wen Teng, Chengxun Xu, Hong Yu, Lingfeng Kong, Shikai Liu, Qi Li

## Abstract

Apart from DNA-sequence-based inheritance, inheritance of epigenetic marks such as DNA methylation is controversial across the tree of life. In mammals, post-fertilization and primordial germ cell reprogramming processes erased most parental DNA methylation information. In nonmammalian vertebrates and insects, it has been proposed that DNA methylation is an essential hereditary carrier. However, how and to what extent general DNA methylation reprogramming affects intergenerational inheritance in molluscs remains unclear. Here, we investigated genome-wide DNA methylation in a mollusc model, the Pacific oyster (*Crassostrea gigas*), to test how epigenetic information transfers from parents to offspring. Analysis of global methylome revealed that the DNA methylation patterns are highly conserved within families. Almost half of the differentially methylated CpG dinucleotides (DMCs) between families in parents could transfer to offspring. These results provided the direct evidence for the hypothesis that the Pacific oyster DNA methylation patterns are inherited in generations. Moreover, distinct DNA methylation differences between male and female somatic tissues in *C. gigas* are revealed in this study. These sex-differential methylated genes significantly enriched in the regulation of Rho protein signal transduction process, which indicated that DNA methylation might have an essential role in the sexual differentiation of somatic tissues in *C. gigas*.

**Author Summary:** Transgenerational inheritance of DNA methylation marks varies across the tree of life. In mammals, post-fertilization and primordial germ cell reprogramming processes obstructed the DNA methylation transmission from parents to child, and only some CpG dinucleotides retain gamete-inherited methylation. However, the DNA methylation inheritance seems apparent in nonmammalian vertebrates and insects. As one of the essential mollusc models, the Pacific oyster *Crassostrea gigas* have received the most substantial epigenetic studies, mainly focusing on the DNA methylation profiles. While a previous study suggested the existence of paternal inheritance of DNA methylation patterns in *C. gigas*, more data are needed to confirm this hypothesis. In this study, genome-wide DNA methylation analysis was performed to investigate the epigenetic inheritance in *C. gigas*. Almost half of the DNA methylation differences between families in parents were found to be transferred to children, indicating the absence of global DNA methylation reprogramming in *C. giga*s. Besides, extensive hypomethylation in *C. giga*s females compared with males were also unveiled. These hypomethylated genes were significantly enriched in the regulation of Rho protein signal transduction process. For example, guanine nucleotide exchange factors, including *KALRN, FGD1*, and *FGD6*, were hypomethylated in *C. gigas* females, and the corresponding transcriptions were significantly upregulated. Our findings provided insights into the evolution of DNA methylation patterns, transgenerational epigenetic inheritance, and sexual differentiation in molluscs.

## Introduction

DNA methylation is a universal epigenetic regulatory mechanism found in prokaryotes and eukaryotes [1, 2]. This prevalent epigenetic modification is essential for bacteria restriction-modification (RM) systems [3] and mammal immune response [4], insect social behavior, embryonic development [5], genome imprinting [6], inactivation of X-chromosome and tissue-specific functions [7, 8]. In mammalian genomes, methylation usually occurs in CpG dinucleotides, with more than 70% of CpG sites methylated [9]. Unlike the densely methylated genome in vertebrates, genomic DNA methylation levels in invertebrates vary across taxa. In insects, Diptera (fruit fly) nearly lost DNA methylation because of the absence of DNA methyltransferase homologs. Hymenopteran (ants, bees, and wasps) had less than 4% DNA methylation levels, while Blattodea had relatively higher levels of genomic DNA methylation, ranging from 1% to 14% [10]. Similarly, cytosine methylation most occurs in CpG dinucleotides and the whole-genome DNA methylation levels are highly variable in Molluscs ranging from 5 to ∼15% [11].

Genetic information provides the primary substrate of inheritable traits across generations. Apart from the DNA sequence-based inheritances, many phenomena of epigenetic-based inheritances, including DNA methylation, histone modification, and small non-coding RNAs, have been reported in organisms [12, 13]. In mammals, genome-wide DNA methylation traits experienced erasure and establishment twice: first after the fertilization and second during the germ cell formation [5]. Two waves of DNA methylation reprogramming are the obstruction of DNA methylation inheritance. Only a small number of parentally imprinted genes escaped reprogramming in the early development of the embryo [14, 15]. However, nonmammalian vertebrates do not undergo genome-wide DNA methylation reprogramming during embryogenesis [16]. For example, zebrafish retained the paternal epigenetic memory in primordial germ cells (PGC) in stark contrast to the findings in mammals [17, 18]. In invertebrates, limited studies give details of the DNA methylation remodeling and epigenetic inheritance. Recent investigations revealed that DNA methylation reprogramming during embryogenesis was absent in cnidarians and protostomes such as insects [16]. For instance, honey bees are reported to have highly conserved DNA methylation patterns between generations [19]. Stable inheritance of an epigenetic signal in *Nasonia* was also found in F_1_ hybrids [20]. These results suggested that the DNA methylation reprogramming seems to be a mammalian-specific feature [19].

As a typical Mollusc model species, the Pacific oyster *Crassostrea gigas* has moderate genomic DNA methylation levels in CpG dinucleotides, ranging from 12 to 18% due to sampling status and methylation calling methods in various studies. Because of the ecological and economic values, *C. gigas* has the most extensive DNA methylation studies in Molluscs [11]. These works are primarily concerned with gene expression regulation [21-23], development processes [24, 25], phylotypic plasticity [26, 27]. While the previous study hypothesized that intergenerational inheritance in DNA methylation exists in *C. gigas* [25], more evidence is needed to make general collusions. Moreover, among most of the epigenetic studies in *C. gigas*, the sex differences in somatic tissues were neglected. One reason is the difficulty in gender determination out of spawning season. The other reason is underestimated genome-wide DNA methylation differences between male and female somatic tissues in *C. gigas*.

Here, we produced diploid and triploid Pacific oysters in two independent families. Whole-genome bisulfite sequencing (WGBS) of both parent and offspring muscle tissues were then performed in each family to investigate the epigenetic inheritance, sexual differences, and effects of chromosome ploidy in DNA methylation in *C. gigas*. We wish our work could add one more puzzle piece to the image of epigenetic studies in Molluscs.

## Results

### Globally DNA methylation landscape of *C. gigas*

To profile the inheritance patterns of DNA methylation in *C. gigas*, regular F_1_ diploid and triploid oysters were produced by crossing a normal diploid male and female oysters in two independent families. The whole-genome bisulfite sequencing (WGBS) was then performed using muscle tissues from both parents, three diploid and three triploid offspring individuals in each family. RNA sequencing (RNA-seq) was also introduced to profile the transcription in the same muscle tissues in offspring (S1A Fig). In total, an average of 66.8 million 150 pair-end reads covering 9 million CpGs (> 68% of the total CpG sites) at least five times were obtained in each sample (S1 Table). DNA methylation ratio was relatively consistent with increasing read depth, which excluded the sequencing depth-induced bias in methylation calling (S1B Fig). Bisulfite conversion efficiencies reached 99.9% in all analyzed samples. The average DNA methylation levels ranged from 0.11 to 0.14 (Fig 1A), consistent with previous studies [26, 27]. Compared with public WGBS data of *C. gigas* [21, 22, 26, 27]. high Pearson correlation coefficients (average *r* = 0.858) were found between our sequencing data and public datasets (S1C Fig). In all, the WGBS data quality in this study is solid.

**Fig 1.**
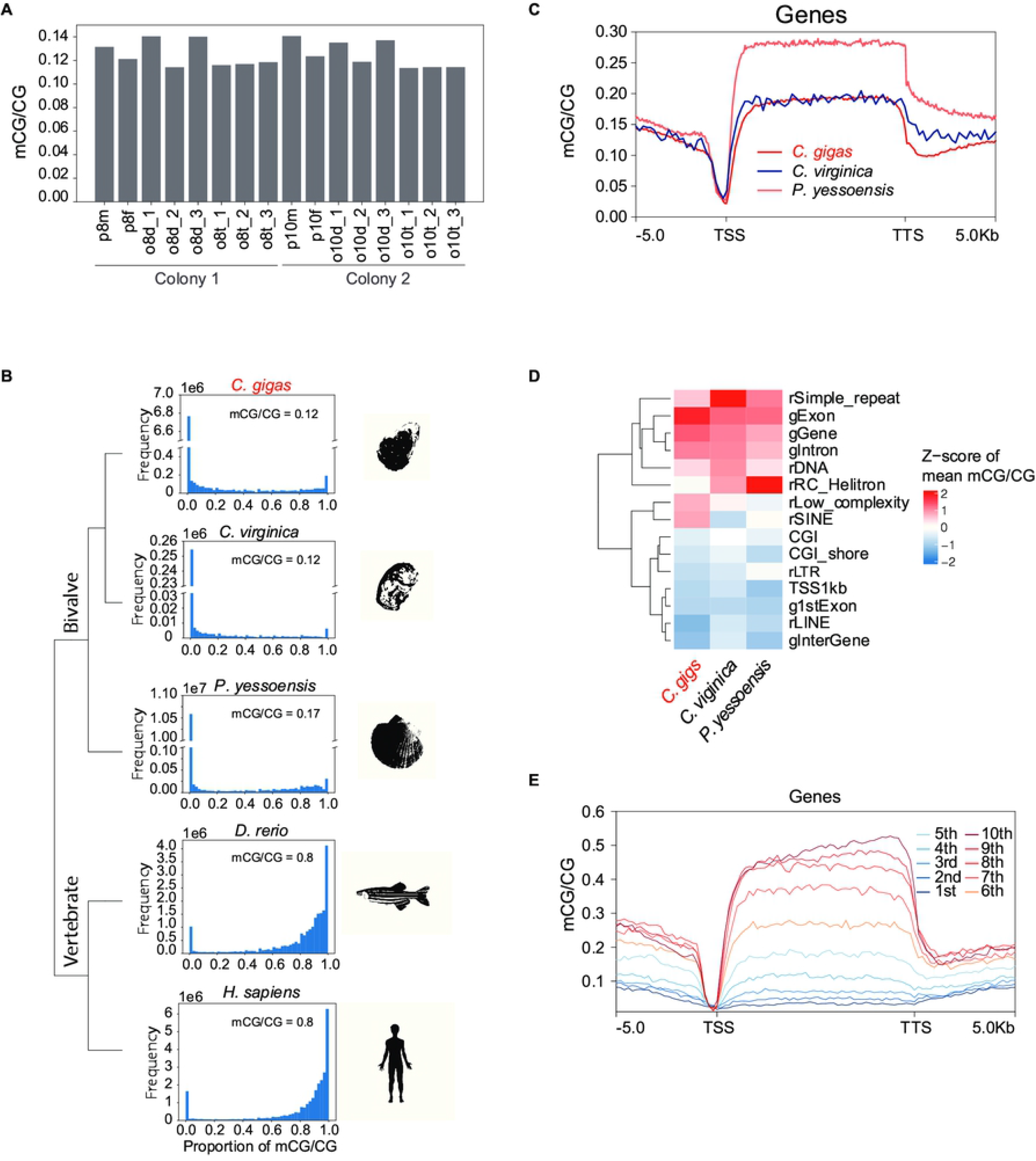
DNA methylation profiles of *C. gigas* muscle tissues. (A) Global DNA methylation levels (quantified as mean mCG/CG) of oysters ranged from 0.11 to 0.14. (B) Histograms of DNA methylation levels distributions for human (*Homo sapiens*) muscle cells, zebrafish (*Danio rerio*) muscle cells, Yesso scallop (*Patinopecten yessoensis*) mantle tissues, eastern oyster (*Crassostrea virginica*) reproductive tissues, and the Pacific oyster (*C. gigas*) muscle tissues in this study. (C) DNA methylation levels across the gene body in *C. gigas, C. virginica*, and *P. yessoensis*. (D) DNA methylation levels at the indicated regulatory elements in *C. gigas, C. virginica*, and *P. yessoensis*. (E) The relationship between DNA methylation levels and transcripts across the gene body.

Next, the *C. gigas* DNA methylation profile was compared with that of two other bivalve model organisms, *Crassostrea virginica* [28] and *Patinopecten yessoensis* [26], to investigate the conserved and derived DNA methylation patterns in bivalves. Under the same data analysis criteria, the average DNA methylation level of *C. gigas* (mCG/CG = 0.12) was consistent with that of *C. virginica* (mCG/CG = 0.12) but relatively lower than *P. yessoensis* (mCG/CG = 0.17). The frequency of DNA methylation ratios of *C. gigas, C. virginica*, and *P. yessoensis* displayed a non-classical bimodal distribution with a major peak at 0 (unmethylated) and a minor peak at 1 (fully methylated), which is distinct from that of vertebrates (Fig 1B and S1D Fig). However, hypermethylated gene body and hypomethylated transcriptional start sites (Fig 1C and S1E Fig) are like other eukaryotes [29]. To further claim the DNA methylation patterns in bivalves, we compared the DNA methylation levels within various regulatory elements. We observed consistently high methylation levels in bivalve genomic regions, including exon, intron, simple repeats, and DNA transposons, but low methylation in CpG islands (CGIs), promoters, long interspersed nuclear elements (LINEs), and long terminal repeats (LTRs). Despite these consistencies, there were low DNA methylation levels within rolling-circle transposons but relatively high DNA methylation levels in low complexity repeats and short interspersed nuclear elements (SINEs) in *C. gigas* (Fig 1D and S1F Fig).

The function of DNA methylation to repress transcription has long been recognized [30]. In *C. gigas*, TSS regions remained almost absent of DNA methylation in both active and inactive genes and showed no strict linear inverse relationship with the transcription (Fig 1E and S1G Fig). The gene body methylation plateau is reported to exhibit a parabolic relationship with transcription: moderately expressed genes are most likely to be methylated, whereas the most active and non-active genes have lower methylation levels [21, 26]. However, our data showed that gene body methylation in both diploid and triploid oysters had a linear relationship with gene expression (Fig 1E and S1G Fig).

### DNA methylation inheritance and sex-specific DNA methylation differences in *C. gigas*

The gender of parent cohort was determined by checking germ cells during the breeding process. Because progenies were out of sexual maturity stages, offspring samples were sexed by DNA methylation markers at the diacylglycerol kinase delta (*DGKD*) locus, which was hypermethylated in males and hypomethylated in females [31]. The DNA methylation marker was reconfirmed in parent groups and public data (S2A Fig). All triploid and two diploid samples were found to be females, while the other four diploid oysters were males (S2B Fig and S2 Table).

Next, we compared the methylation density using all sample shared CpGs in each colony (excluded common CpGs with zero methylation level in all specimens) to test whether there were global changes of DNA methylation between generations and different genders. Overall, parents and progenies had similar methylation densities, but female DNA methylation levels were found to be lower than males (*P* < 0.00001; Fig 2A). These hypomethylation patterns in females spread across chromosomes and enriched in genomic region including exon, intron, and low complexity regions (S2C Fig). Unsupervised hierarchical clustering analysis based on the top 10,000 variable common CpGs across all samples distinguished females from male groups. Under the main clusters of different genders, samples from colony one constituted a distinct sub-cluster, separated from colony two (Fig 2B). These findings revealed distinct global DNA methylation differences across genders and families in *C. gigas*.

**Fig 2.**
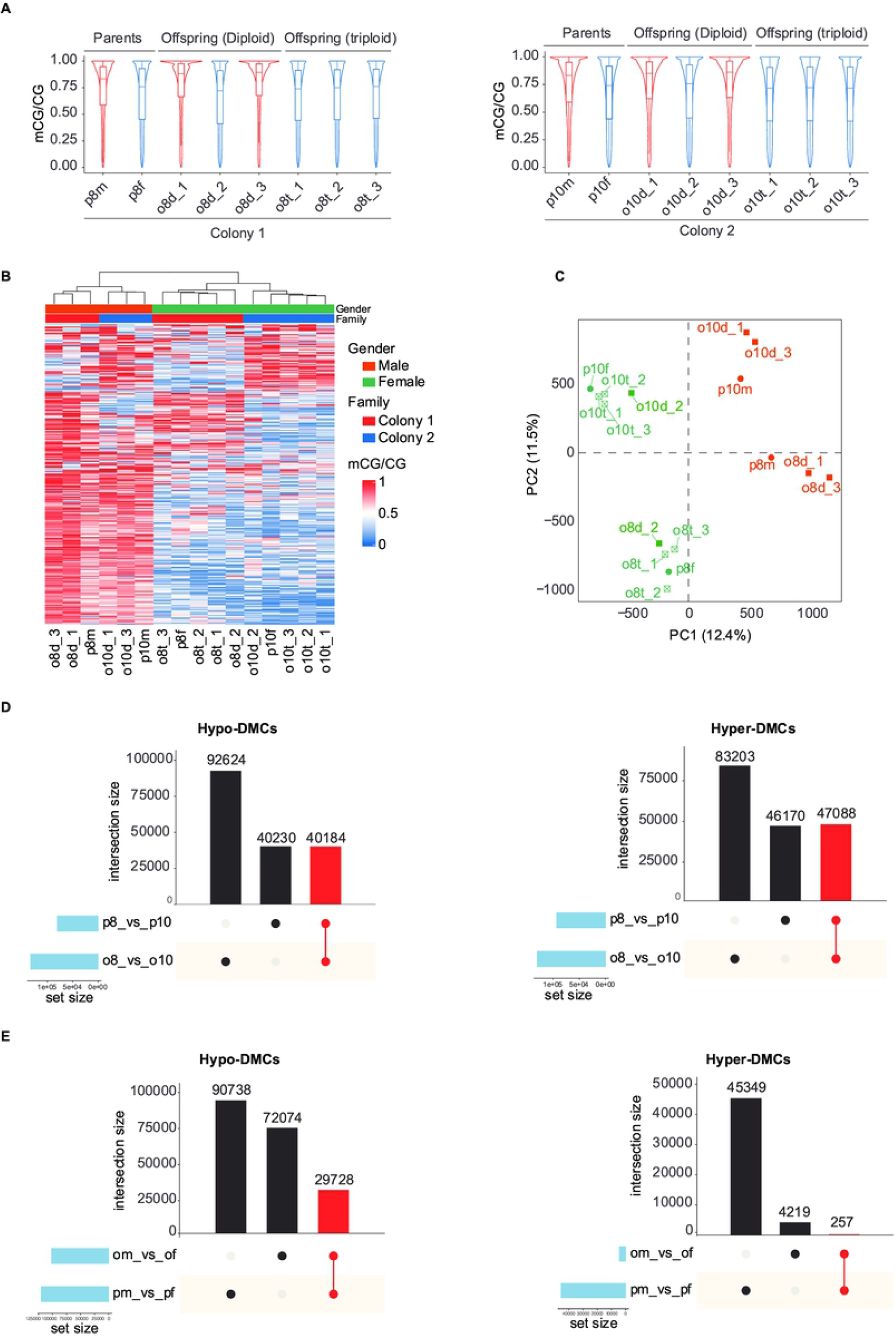
Inheritance and sex differences of DNA methylation in *C. gigas* muscle tissues. (A) DNA methylation levels across samples in colony one (left) and two (right). Violin plots represent kernel density plot. Boxplots represent median and interquartile range. (B) Heatmap of DNA methylation levels using the top 10,000 variable common CpGs across all samples in this study. (C) Principal component analysis (PCA) of DNA methylation levels in shared CpGs across all samples in this study. (D) UpSet plots showing the integrated comparative analysis of hypo-DMCs (left) and hyper-DMCs (right) between parents and offspring. DMCs are identified between colonies. (E) UpSet plots showing the integrated comparative analysis of hypo-DMCs (left) and hyper-DMCs (right) between parents and offspring. DMCs are identified between males and females.

To confirm these findings, Pearson correlation analysis was performed using pairwise common CpGs across all samples analyzed in this study. There were relatively high Pearson correlation coefficients among samples in two main clusters (male and female clusters) and even higher Pearson correlation coefficients among samples in colony subclusters (S2B Fig). Principal component analysis using all sample shared CpGs, including 5.9 million CpG dinucleotides was also conducted. The principal component one divided all individuals into male and female groups. Meanwhile, the principal component two divided all samples into two colony groups (Fig 2C).

To evaluate whether differences between colonies, male and female groups in parents were recapitulated in offspring. We identified differentially methylated cytosines (DMCs) by comparing colony one and colony two, male and female groups in parents and offspring using MOABS [32], respectively. In progenies, we found a great deal of DMCs between colonies, male and female groups, but much fewer DMCs between diploid and triploid groups (S2C Fig). These results reconfirmed that there are no global DNA methylation differences between diploid and triploid oysters [33]. Furthermore, we found almost half of the DMCs, including hypo-DMCs and hyper-DMCs, between colony one and colony two in parents transferred to offspring (Fig 2D). Moreover, female groups in offspring displayed a global decrease in DNA methylation, consistent with that of parents (S2E Fig). And a great deal of hypo-DMCs between males and females in parent cohorts were recapitulated in offspring (Fig 2E).

Overall, our results indicated that there is no DNA methylation reprogramming in *C. gigas*, and almost half of the family-specific DNA methylation marks could be stably transferred between generations. Besides, sexual differentiation in DNA methylation profiles exist in *C. gigas* somatic tissues.

### Activation of Rho signaling in *C. gigas* females

To investigate DNA methylation differences between males and females in *C. gigas* muscle tissues, the differentially methylated regions (DMRs) were identified by comparing the male and female groups in parents and offspring, respectively. In consequence, 10,180 hypo-DMRs and 3,555 hyper-DMRs were found in parent cohort. Similarly, 9,794 hypo-DMRs and 196 hyper-DMRs were found in offspring cohort (Fig 3A). Across generations, 4,148 hypo-DMRs and 23 hyper-DMRs were shared by parent and offspring cohorts (Fig 3B), which suggested a stable decrease of DNA methylation in females in *C. gigas* muscle tissues. These stably inherited hypo-DMRs in females scattered across all chromosomes and mainly enriched in gene bodies, especially in intron regions (S3A and B Fig). AgriGO v2.0 [34] was then used to perform gene ontology (GO) enrichment analyses. We found these common hypo-DMR related genes are highly enriched in the regulation of Ras homology (Rho) protein signal transduction process (Fig 2C and S3C Fig). The Rho GTPases switch cycled between the inactive (GDP-bound) and active (GTP-bound) forms, regulated by guanine nucleotide exchange factors (GEFs) and GTPase-activating proteins (GAPs) (Fig 3D). Active GTP-bound GTPases interact with various downstream effectors and regulate a wide range of cellular responses [35]. In this study, 21 genes function as GEFs were found to be hypomethylated in females compared to males (S3 Table). For example, the DNA methylation levels at *KALRN, FGD1*, and *FGD6* locus were significantly decreased in females compared with that in males in *C. gigas* (Fig 3E and S3D Fig). Hypomethylation consequently promoted the gene expression as the transcriptions of *KALRN, FGD1*, and *FGD6* of females were significantly higher than that of males in *C. gigas* (Fig 3F and S3E Fig).

**Fig 3.**
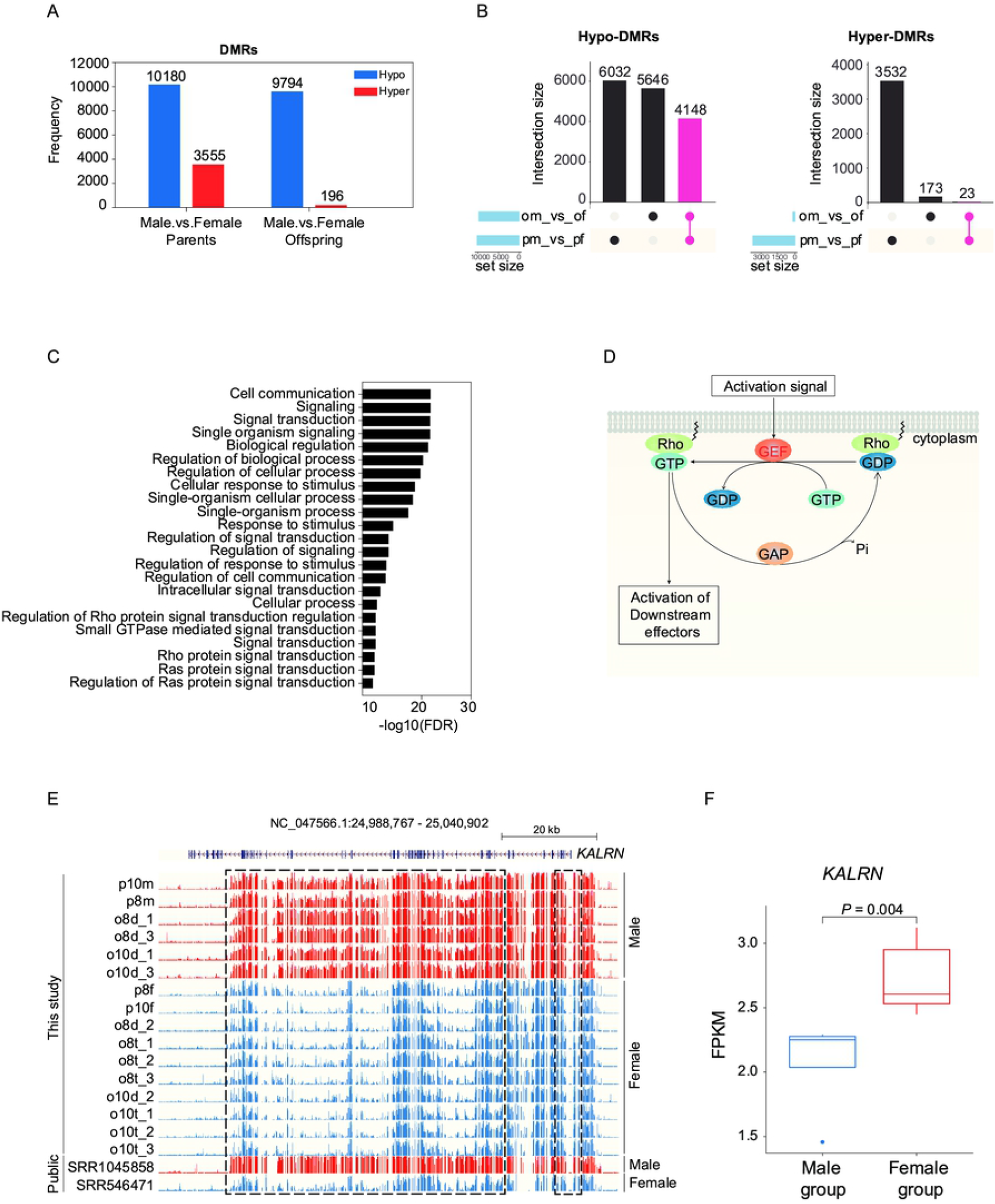
Functional annotation of cytosine methylation differences between male and female in *C. gigas*. (A) Histogram showing the numbers of hypo-DMRs and hyper DMRs identified between males and females in parent groups and offspring groups. (B) UpSet plots showing the integrated comparative analysis of hypo-DMRs (left) and hyper-DMRs (right) between parents and offspring. DMRs are identified between males and females in *C. gigas*. (C) Gene ontology (GO) enrichment analysis for shared hypo-DMRs related genes across parent and offspring cohorts. Hypo-DMRs are identified between males and females in *C. gigas*. The x-axis shows the false discover rate (FDR) value. (D) Overview of Rho GTPase regulation. (E) UCSC genome browser view of DNA methylation enrichment at the *KALRN* locus (NC_047565.1:11384273-11464307) in all samples analyzed in this study. The highlighted region by dotted line exhibits decreased methylation in females. (F) Boxplot showing the FPKM values in male and female groups in offspring (*P* < 0.01).

These results suggested that epigenetic differences exist between males and females in *C. gigas* muscle cells, and the DNA methylation signature at GEFs genes can be used as biomarkers to distinguish the gender of *C. gigas*. Besides, these intrinsic epigenetic differences in the Rho protein signal transduction process may contribute to sex-dependent differences in *C. gigas* muscle phenotypes.

## Discussion

DNA methylation is prevalent in eukaryotic organisms [36]. This epigenetic modification mechanism is especially predominant in vertebrates but varies greatly in invertebrates. In bivalves, the global DNA methylation levels of CpG dinucleotides have been shown conventional invertebrate-like patterns with a majority CpGs unmethylation [11]. However, the unmethylated CpG islands and apparent gene-body methylation in bivalves were like that of other eukaryotic organisms; these conserved methylation patterns may serve as an ancient feature through the evolution of eukaryotes [29, 37]. In oyster and scallop genomes, we found apparent DNA methylation in some repetitive elements, including simple repeat and DNA transposon. In contrast, the methylation of some retrotransposons, including LTR and LINE, occurs only at moderate levels. Despite these consistencies, differences in cytosine methylation also exist within bivalve species. For example, the global cytosine methylation in scallops is higher than in oysters. *C. gigas* also showed some species-specific methylation patterns. The rolling-circle transposable elements, *helitrons*, in *C. gigas* displayed a moderate methylation level. *Helitrons* amplified significantly in *C. gigas* genome and were proposed to be remnants of the past activity of evolution [38]. The diminish of cytosine methylation in *Helitrons* may also be the result of ancient activations, as deamination is often needed for transposable elements to take on regulatory functions [39].

The repressive effect of DNA methylation at promoters on transcription initiation has long been recognized [30]. High methylation levels at promoters may exclude the DNA-binding factors and consequently depress the transcription [40]. However, gene body methylation is positively correlated with gene expression [41, 42]. It was proposed that the DNA methylation in the gene body facilitates the transcription elongation and affects splicing [43, 44], and that it inhibits intragenic promoters [45]. In *C. gigas*, DNA methylation is predominantly enriched in intragenic regions, especially in exon. Similarly, this gene-body methylation is positively correlated with transcriptions. These conservative patterns and functions indicated the fundamental roles of DNA methylation in *C. gigas*.

The absence of DNA reprogramming has been observed in cnidarians and protostomes [16]. For example, DNA methylation marks are stably transferred between generations in honey bees [19]. Besides, a previously underestimated fraction of the vertebrate genome could even bypass the DNA methylation reprogramming process [13, 46]. However, the epigenetic inheritance in molluscs remains poorly understood. A previous study had suggested that DNA methylation patterns are inherited in *C. gigas* [25], but direct evidence for this hypothesis was no longer provided. Our data corroborated that almost half of the methylation differences between colonies in parents could transfer to the next generations. The stable DNA methylation inheritance in *C. gigas* provides the basis to study the environmentally induced epigenetic changes and inheritance.

DNA methylation has been reported to differ males and females in mammalian tissues, including islets [47], brain [48], and skeletal muscle [49]. However, most mollusc studies neglected these differences between genders, especially in *C. gigas*. Researchers consistently underestimated the differences between males and females in *C. gigas* somatic tissues. In this study, distinct DNA methylation profiles between male and female muscle tissues were unveiled in *C. gigas* (Fig 3A and B). These epigenetic differences (DMRs) are not enriched in solo chromosomes but scattered across the genome (S3A Fig). Besides, the DMRs between males and females are enriched in genetic regions, especially in intro regions, indicating the potential genetic regulation roles of the methylation alteration. Gene ontology analyses revealed significant enrichment for the regulation of Rho protein signal transduction process across the differentially methylated genes. Specifically, 21 GEFs genes activating the GTPase by exchanging bound GDP for free GTP were hypomethylation in females. The transcription of three GEFs, including *KALRN, FGD1*, and *FGD6*, were also found upregulated in females compared with males. Rho GTPases are highly conserved across all eukaryotes and are best known for their roles in several cellular processes, including cytoskeletal organization, cell cycle progression, apoptosis, and membrane traffic [35]. Previous studies have shown Rho GTPases have a critical role in human muscle development, regeneration, and function [50, 51]. The distinct nucleotide methylation differences at these GEFs locus between *C. gigas* males and females indicated that Rho GTPase signaling might contribute to the muscular phenotypes. Therefore, we highly recommend taking gender into consideration in epigenetic studies in *C. gigas*.

DNA methylation is an essential epigenetic modification mechanism, and it has long been shown involved in *C. gigas* gene expression, embryonic development, growth, sex differentiation, genetic inheritance, and phenotype plasticity. This study emphasized the influences of DNA methylation marks in genetic inheritance and sexual differentiation in somatic tissue developments in *C. gigas*. But so far, we still lack large pieces of the entire DNA methylation landscapes of *C. gigas*. For example, tissue-specific DNA methylation patterns, gametogenesis and embryo development DNA methylation dynamics at base resolution are poorly understood. Future work towards these basic epigenetic studies in *C. gigas* is required.

## Conclusion

The present work provided direct evidence that DNA methylation patterns could transfer between generations. We hypotheses that there is no global DNA methylation reprogramming in *C. gigas*. Distinct DNA methylation differences exist between male and female oyster somatic tissues. The CpG dinucleotides alteration in Rho GTPases cycle may control the sex-based differences in muscular phenotypes. Specifically, hypomethylation in GEFs in *C. gigas* females activates the Rho GTPases switch and activates the downstream factors. These findings provide new insights into the DNA methylation influences in genetic inheritance and sexual differentiation in molluscs.

## Materials and methods

### Animals

One normal diploid male and female oysters were selected for mating from two full-sib families, respectively. In each family, fertilized eggs were divided into two equal groups. One group was treated with cytochalasin B (CB, 0.5 mg L−^1^) for 15 min once 50 % of the eggs released the first polar body to produce the triploid oysters. The other untreated group produced diploid oysters normally. Progenies were reared separately for one year. Chromosome ploidy of each sample was determined using flow cytometry to check the whole genome DNA contents stained by DAPI. In each family, adductor muscles from male and female parents, three diploid progenies, and three triploid progenies were frozen by liquid nitrogen and then transferred to a -80 °C refrigerator for long-term preservation.

### WGBS library construction and data analysis

Genomic DNA (gDNA) was isolated from adductor muscle tissues in parents and offspring using the TIANamp Marine Animals DNA Kit (TIANGEN, Beijing). Library preparation and high-throughput sequencing were conducted by Novogene (Beijing, China). Briefly, approximately 5.2 µg of purified gDNA (spiked with 1% unmethylated lambda DNA, Promega) was sheared into fragment size of 200-300 bp using Covaris S220. These DNA fragments were then subjected to bisulfite conversion using EZ DNA Methylation-GoldTM Kit (Zymo Research). The resulting bisulfite-converted DNA fragments were amplified by PCR and then purified by AMPure XP beads (Beckman Coulter). Finally, the library was sequenced on Illumina Hiseq platform with cBot System via TruSeq PE (Paired-End) Cluster Kit v3-cBot-HS (Illumina, US). For WGBS data analysis, raw FASTQ data were filtered using *fastp* v.0.20.1 [52] with main parameters (--cut_front --cut_front_window_size=1 --cut_front_mean_quality=3 --cut_tail --cut_tail_window_size=1 --cut_tail_mean_quality=3 --cut_right --cut_right_window_size=4 --cut_right_mean_quality=15 --trim_front1 10 --trim_front2 10). The filtered FASTQ files were then mapped to *C. gigas* reference genome (GCF_902806644.1) using bsmap v.2.90 [53] with parameters (-R -p 4 -n 1 -r 0 -v 0.1 -S 1). BSeQC [54] were used to evaluate the quality of bisulfite sequencing output. The CpG coverage and DNA methylation calling were performed using MCALL module in MOABS v.1.3.0 [32]. In this study, only CpG sites sequenced by at least 5 times were retain in the following analysis. Bisulfite conversion efficiency was estimated by spike-in unmethylated lambda DNA. The significant differentially methylated CpG sites (DMCs) and differentially methylated regions (DMRs) were identified using MCOMP module in MOABS with main parameter (--withVariance 1). Bigwig files were upload to UCSC genome browser for visualization. Functional annotations were performed using AgriGO v2.0 [34].

### mRNA-seq library construction and data analysis

Total RNA was isolated from muscle tissues in offspring using RNAprep Pure Tissue Kit (TIANGEN, Beijing). According to the manufacturer’s instructions, library constructions were performed using TruSeq Stranded mRNA LT Sample Prep Kit (Illumina, US). In brief, samples with the RNA integrity number (RIN) ≥ 7 were used in the following two rounds of mRNA purification using oligo-dT beads to capture polyA tails. RNA fragmentations were performed by Covaris S220. Then the first strand cDNA was synthesized by reverse transcribing the cleaved RNA fragments primed with random primer. Second strand cDNA was synthesized via incorporating dUTP in place of dTTP, and the dUTP strand degraded in the following amplification process. One adenine nucleotide was added to the 3’ ends of blunt fragments. Finally, indexing adapters were ligated to the ends of the double-strand cDNA fragments. Each library was deeply sequenced on Illumina NovaSeq 6000.

For RNA-seq data analysis, raw FASTQ data was filtered using *fastp* v.0.20.1 as previously described. The filtered FASTQ files were then mapped to *C. gigas* reference genome (GCF_902806644.1) using HISAT2 v.2.2.1 [55]. Mapped reads with mapping quality ≥ 30 were retained in the following analysis. Read counts and FPKM was calculated using HTSeq 2.0 [56].

### Public data

Public WGBS data of 22 *C. gigas* samples were downloaded from PRJNA213124, PRJNA173440, PRJNA562805, and PRJNA689936. WGBS data of four *H. sapiens* samples were downloaded from PRJNA63443. WGBS data of four *D. rerio* samples were downloaded from PRJNA553572 and PRJNA628650. WGBS data of six *P. yessoensis* samples were downloaded from PRJNA695315. Reduced representation bisulfite sequencing (RRBS) data of 77 *C. virginica* samples were downloaded from PRJNA488288.

All public data were analyzed under the same criteria as described above. *H. sapiens* reads were mapped to NCBI Human Reference Genome Build GRCh38 (hg38). *D. rerio* reads were mapped to GCF_000002035.6 (GRCz11). *P. yessoensis* reads were mapped to GCF_002113885.1 (ASM211388v2). *C. virginica* reads were mapped to GCF_002022765.2 (C_virginica-3.0). Considering the low coverage of RRBS data of *C. virginica*, we merged all data into one sample. Transposable element files of *H. sapiens* and *D. rerio* were downloaded from NCBI. Putative transposable elements of *P. yessoensis, C. virginica*, and *C. gigas* were identified with RepeatMasker v4.1.2 [57] using the mollusca RepBase repeat library [58] and RepeatModeler [59].

## Data availability

WGBS and RNA-seq data are available at the NCBI under the project number PRJNA801419. All relevant data supporting our findings are available within the article and supplementary information files or from the corresponding author for reasonable request.

## Acknowledgments

This work was supported by the China Agriculture Research System Project (CARS-49), and Earmarked Fund for Agriculture Seed Improvement Project of Shandong Province (2020LZGC016).

## Supporting information

**S1 Fig. Standard quality control of DNA methylation analysis in *C. gigas***

(A) Schematic of the experimental design. Two pairs of parents (n = 4), diploid oysters (n = 6) and triploid oysters (n = 6), were used in this study. (B) The relationship between DNA methylation levels and sequencing depth. And the relationship between coverage of CpG sites and sequencing depth. (C) Pearson correlation analysis between WGBS data in this study and public WGBS data of *C. gigas*. (D) DNA methylation levels across the gene body of all *C. gigas* samples analyzed in this study. (E) Histograms of DNA methylation levels distributions for all analyzed samples. (F) DNA methylation levels at the indicated regulatory elements in all analyzed samples. (G) The relationship between DNA methylation levels and transcripts across the gene body in triploid oyster samples.

**S2 Fig. DNA methylation differences between colonies, male and female, and diploid and triploid oysters**

(A-B) The University of California, Santa Cruz (UCSC) genome browser view of DNA methylation enrichment at the *DGKD* locus (NC_047562.1:45,716,336-45,757,148) in parent cohort (A) and offspring cohort (B). The highlighted region exhibits decreased methylation in females. (C) Heatmap of the DNA methylation levels in regulatory elements across all samples analyzed in this study. (D) Heatmap of the Pearson correlation coefficient using pairwise common CpGs methylation levels across all samples. (E) Histogram showing the numbers of hypo-DMCs and hyper DMCs identified between colony one and two, males and females, diploid and triploid oysters.

**S3 Fig. Differentially methylated genes between male and female in *C. gigas***

(A) Histogram showing the distribution of hypo-DMRs in chromosomes. (B) Histogram showing the distribution of hypo-DMRs in regulatory elements. (C) The hierarchical structure and ancestry relationships in the gene ontology top enriched terms. (D) UCSC genome browser view of DNA methylation enrichment at the *FGD1* locus (NC_047566.1:24989612-25017157) (left) and *FGD6* locus (NC_047566.1:21330721-21358510) (right) in all samples analyzed in this study. The highlighted region by dotted line exhibits decreased methylation in females. (E) Boxplot shows FPKM values of *FGD1* (left) and *FGD6* (right) in male and female groups in *C. gigas* (*P* < 0.05).

**S1 Table. Sample information and WGBS data statistics**.

16 samples collected from two independent families included parents and progenies. 2×75bp paired-end sequencing was performed in this study. The results of WGBS data analysis included numbers of filtered FASTQ reads (Total reads), unique mapped reads (Uniq mapped reads), unique mapping ratio (Mapping ratio), bisulfite conversion efficiencies estimated from spike-in Lambda genome (Bcr Lambda), bisulfite conversion efficiencies estimated from whole genome (Bcr whole genome), CpG sites with at least 5 times coverage (Number of CpGs), effective reads ratio (Positive rate), global CpGs methylation ratio (Mean ratio of CpGs).

**S2 Table. Results of gender determination across all samples analyzed in this study**.

Males included p8m, o8d_1, o8d_3, p10m, o10d_1, o10d_3. Females included p8f, o8d_2, o8t_1, o8t_2, o8t_3, p10f, o10d_2, o10t_1, o10t_2, o10t_3.

**S3 Table. Regulation of Ras protein signal transduction process enriched in Gene ontology analysis**.

In regulation of Ras protein signal transduction process significantly enriched in this study, 21 genes functioning as guanine nucleotide exchange factors (GEFs) were hypomethylated in *C. gigas* females.

